# Toxicokinetics of citreoviridin *in vivo* and *in vitro*

**DOI:** 10.1101/578302

**Authors:** Yosuke Uchiyama, Masahiko Takino, Michiko Noguchi, Nozomi Shiratori, Naoki Kobayashi, Yoshiko Sugita-Konishi

## Abstract

Citreoviridin (CIT) produced by *Penicillium citreonigrum* as a secondary metabolite is a yellow rice toxin that has been reported to be related to acute cardiac beriberi; however, its toxicokinetics remain unclear. The present study elucidated the toxicokinetics through swine *in vivo* experiments and predicted the human toxicokinetics by a comparison with findings from *in vitro* experiments. Swine *in vivo* experiments revealed that CIT had a high bioavailability of more than 90%. In addition, it showed a large volume of distribution (1.005 ± 0.195 L/kg) and long elimination half-life (17.7 ± 3.3 h) in intravenous. These results suggested the possibility of a slow metabolism of CIT. An intestinal permeability study using the human cell line Caco-2 showed that CIT had a high permeability coefficient, suggesting it would be easily absorbed in human intestine, similar to its absorption in swine. The metabolite profiles were investigated by incubating CIT with S9 obtained from swine and humans. Hydroxylation, methylation, desaturation and dihydroxylation derivatives were detected as the predominant metabolites, and CIT glucuronide was produced slowly compared with above metabolites. A comparison of the peak area ratios obtained using quadrupole time-of-flight mass spectrometer showed that the rates of all of the main metabolites except for glucuronide produced using human S9 were three-fold higher than those obtained using swine S9. Furthermore, the elimination of CIT using human S9 was more rapid than when using swine S9, indicating that CIT would be metabolized faster in humans than in swine. These *in vivo* results suggested that CIT is easily absorbed in swine and persists in the body for a long duration. Furthermore, the CIT metabolism appeared to be faster in human liver than in swine liver *in vitro*, although the bioavailability of CIT was predicted to be similarly high in humans as in swine.

## Introduction

Citreoviridin (CIT) is a mycotoxin produced by *Penicillium citreonigrum*, *Aspergillus terreus* and *Eupenicillium ochrosalmoneum* as a secondary metabolite [1]. Because CIT is mainly found as a contaminant in rice, it is a serious problem in countries where people consume rice as a staple food.

In 2006, an outbreak of beriberi occurred in Brazil, with a reported 40 of 1207 cases dying [2]. Since *P. citreonigrum* and CIT were detected in rice samples, it was suspected that rice was the causative food of beriberi in the area [3]. CIT contaminating yellow rice has been reported to be related to acute cardiac beriberi (so-called “Shoshin-kakke”) [4]. Uraguchi et al. discovered that an ethanol extract from rice infected with *P. citreo-viride Biourge* (current *P. citreonigrum*) caused symptoms in mice similar to acute cardiac beriberi in humans. In addition, Ueno et al. reported that the isolate from *P. citreo-viride BIourge* which was isolated by Sakabe et al. was chemically identical with CIT [4][5]. Although “Shoshin-kakke” was prevalent in Japan in the 17^th^ and 18^th^ centuries, its occurrence markedly decreased in the early 20^th^ century due to the strengthening of policies to remove moldy rice from the market. Such an epidemiological evidence supported the relation of CIT and cardiac beriberi.

In animal experiments [6] using purified CIT, CIT has been shown to cause fatal adverse effects, with symptoms characterized by ascending paralysis, disturbance of central nervous system and respiratory arrest. The lethal dose 50% (LD_50_) of CIT against mice was reported to be 3.6-11.8 mg/kg subcutaneously and 7.5 mg/kg intraperitoneally [7][8]. When crude extract from yellow rice was given to several mammals via subcutaneous (SC), intraperitoneal (IP) and per os (PO), the typical neurological symptoms mentioned above were observed [9]. In addition, the development of these symptoms in rats occurred earlier with increasing dose [10]. Multiple SC administrations for several months resulted in deaths among rats, even when the dose per day was kept at about 1/100 of the LD_50_ [10]. However, the accumulative property of CIT has not been described in much detail.

While several toxicokinetics studies of CIT have been reported, most were published in the 1980s, and their overall numbers are few. No measurable amount of CIT was reportedly detected in urine, although CIT administered to rats was detected in major organs and feces [11]. Little is known about the bioavailability of CIT.

The present study attempted to elucidate the toxicokinetics and bioavailability of CIT and the production of its metabolites through *in vivo* and *in vitro* studies. First, the kinetics of CIT in plasma were investigated by administering CIT intravenously and orally in swine as an *in vivo* study. Subsequently, to estimate the bioavailability in humans, the intestinal permeability of CIT using Caco-2 cells and the metabolism of CIT and production of its main metabolites using S9 fractions obtained from swine and humans were investigated. We herein report the toxicokinetic characterization of CIT *in vivo* and *in vitro* and discuss the behavior of CIT in the body.

## Materials and Methods

### Reagents

CIT (purity: 82.4%) (Fig. 1) was extracted from *P. citreonigrum* isolated by Shiratori et al. in our laboratory [12] with reference to the method of da Rocha et al. [13]. Caco-2 cell lines were provided by the Division of Pharmacognosy, Phytochemistry, and Narcotics of the National Institute of Health Sciences (Kanagawa, Japan). NADP, G-6-P, Protease inhibitor cocktail, Hanks’ balanced salt solution (HBSS) and HEPES were purchased from Sigma-Aldrich (St. Louis, MO, USA). Uridine-5′-diphosphoglucuronic acid trisodium salt (UDPGA) was obtained from Nacalai Tesque, Inc. (Kyoto, Japan). Alamethicin was obtained from LKT Laboratories, Inc. (St. Paul, MN, USA). Inactivated fetal bovine serum was obtained from Biowest (Nuaillé, France). Dulbecco’s modified Eagle’s medium (DMEM), penicillin and streptomycin were purchased from Invitrogen Japan (Tokyo, Japan). Non-essential amino acids were obtained from MP Bio Science (Derbyshire, UK). Corning^TM^ BioCoat^TM^ Intestinal Epithelium Differentiation Environment Kit was purchased from Corning (NY, USA). Medetomidine hydrochloride and butorphanol tartrate were obtained from Meiji Seika Pharma Co., Ltd. (Tokyo, Japan). Midazolam was obtained from Astellas Pharma Inc. (Tokyo, Japan). Other reagents were purchased from FUJIFILM Wako Pure Chemical Corporation (Osaka, Japan).

**Fig 1.**
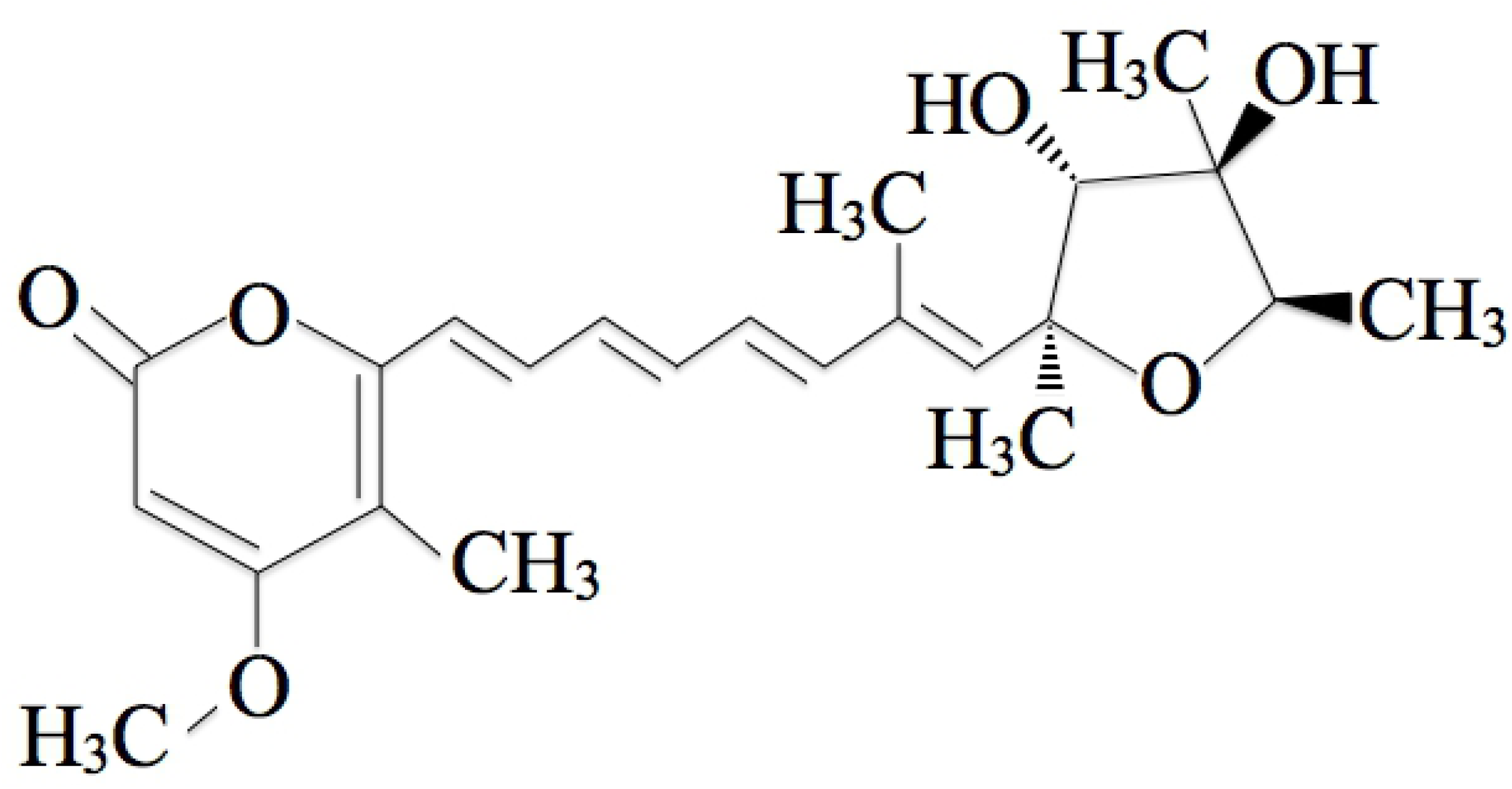
Chemical structure of Citreoviridin.

### S9 fraction preparation

Human hepatic and intestinal S9 fractions and a swine hepatic S9 fraction were purchased from Sekisui XenoTech, LLC. (Kansas City, KS, USA). Swine intestinal S9 fraction was prepared using an intestinal tract of a swine provided by Dr. Shimazu of the Food Physiology Laboratory, Azabu University. Swine intestinal S9 fractions were prepared with reference to the method of Damre et al. [14]. Duodenum specimens were collected from the swine intestinal tract. After washing away the intestinal contents with ice-cold PBS, it was put on an ice-cold tray and the mucosa collected by scratching. Collected mucosa was added 2-3 fold volume of a homogenizing buffer (100 mmol/L potassium phosphate buffer, pH7.4, 150 mmol/L KCl, 250 mmol/L sucrose and 0.1% protease inhibitor cocktail), followed by homogenization. The homogenate was centrifuged at 1000 *g* for 10 min at 4 °C, and the supernatant was transferred to 50-mL centrifuge tubes. The supernatants were centrifuged at 9000 *g* for 20 min at 4 °C to obtain S9 fractions, and each fraction was stored at −80 °C until used in the study.

### Administration study

#### Animals and diets

Swine (barrows; Landrace × Large White × Duroc, 9.4±1.3 kg) were obtained from CIMCO Co., Ltd. (Tokyo, Japan). They were housed in individual cages (0.88 m wide, 1.3 m deep), with free access to water, and fed a commercial feed in quantities of 1.5%-2% of their body weight (BW) daily. All protocols were approved by the Azabu University Ethics Committee of Animal (Approval number: 170829-1).

#### Administration and blood draw

CIT stock solution was prepared by dissolving CIT in acetonitrile to 10 mg/mL. The required amount of CIT was moved from the stock solution into a tube and dried with nitrogen. The dried CIT was then re-dissolved in ethanol:saline (1:4) and used as a test solution. Following three days of acclimatization, administration studies were carried out. CIT (0.1 mg/kg·BW) was intravenously administered to swine (n=4) via the auricular vein. For PO (n=4), 10 mg/mL of CIT-ethanol solution and a small amount of water were added to feed (10 g) in order to make 0.1 mg/kg·BW. This was then fashioned into a sphere and fed to the animals. It was visually confirmed that the animals had eaten the CIT-contaminated feed.

Blood was sampled from the jugular vein at 0 min (before administration), and 5, 10, 20 and 30 min and 1, 2, 3, 4, 8, 24 and 48 h after administration. Blood samples were placed into heparinized tubes and stored on ice until centrifugation. After centrifugation (1919 *g*, 10 min), the plasma was stored at −80 °C temporarily. Plasma samples were prepared according to the method of Devreese et al. [15]. A 3-fold volume of acetonitrile was mixed with the plasma samples, which were then centrifuged again (8500 *g*, 4 °C 10 min) after mixing with a vortex mixer for 15 seconds. The supernatants were transferred into amber screw-top vials and dried under nitrogen gas. The samples were stored at −30 °C until the analysis.

In order to frequently sample blood in a short period of time, up to 1 h after administration, CIT was administered intravenously and orally, followed by immediately administration of 0.1 mg/kg of mixed anesthetics (medetomidine hydrochloride:midazolam:butorphanol tartrate = 3:2:2) via intramuscular injection. Each animal was awake at about one hour after anesthetization.

#### Toxicokinetic analyses

Toxoicokinetics were analyzed with reference to a book [16]. For the toxicokinetic analysis of CIT data obtained via IV, a two-compartment model was applied, whereas a non-compartment model was used for PO data. The bioavailability was determined by calculating the area under the curve (AUC) of IV and PO data, which was determined using the linear trapezoidal rule with extrapolation to infinity.

#### Permeability study using Caco-2 cells

Cell culture and a permeability study were carried out by the method of Kadota et al. [17]. CIT solutions (3 and 10 mmol/mL) were prepared by dissolving dried CIT in DMSO. Permeability study was carried out using Corning^TM^ BioCoat^TM^ Intestinal Epithelium Differentiation Environment Kit (Corning, NY, USA). Cell incubation and induction of differentiation were performed in accordance with the protocol of the kit. CIT solutions were added to Enterocyte Differentiation Medium (EDM) containing 0.08% MITO + serum extender, with CIT concentrations of 3 and 10 µmol/L.

EDM containing CIT was exposed to Caco-2 cells from the apical (AP) side. The transepithelial electrical resistance (TEER) of the AP and basolateral (BL) sides was measured at 0 (before exposure), 1 and 2 h (after exposure) using a Millicell ERS device (Millipore, Molsheim, France). To determine the CIT concentration at the AP and BL sides, transport buffer was collected from both sides. After 400 µL of collected buffer (per side) was transferred to a micro tube, a three-fold volume of acetonitrile was added. Samples were mixed with a vortex mixed, followed by centrifugation at 8500 *g* for 10 min at 4 °C. The supernatant was then transferred to an amber vial and dried with nitrogen gas. Samples were stored at −30 °C until analyzed by liquid chromatography tandem mass spectrometry (LC-MS/MS).

#### Production of CIT metabolites by incubating with S9 fractions

S9 incubation without UDPGA was carried out with reference to a previous report [18]. CIT stock solution (25 µL) was transferred into a micro tube and dried with nitrogen. CIT solution (250 µg/mL) was prepared by re-dissolving dried CIT with 1 mL of DMSO. CIT additive solution (150 µg/mL) was prepared by mixing 600 µL of CIT solution (250 µg/mL) and 400 µL of a base buffer. The total volume of the test solution was 500 µL. The final concentrations of each factor in the test solution were 5 mmol/L MgCl_2_, 5 mmol/L Glucose-6-phosphate and 0.5 mmol/L NADP. The S9 and CIT concentrations in the test solution were 0.5 mg/mL and 1.5 µg/mL, respectively. After adding CIT, the test solution was incubated in a warm bath at 37 °C for 30, 60 and 240 min. The reaction of the test solution was terminated by adding the same amount (500 µL) of acetonitrile. Each sample was mixed in a vortex mixer at 30 s, followed by centrifugation at 6000 *g* for 10 min at 4 °C. The supernatant was transferred to an amber vial and dried with nitrogen gas. Dried samples were stored at −30 °C until the analysis.

S9 incubation with UDPGA was carried out as follows: First, a mixture (S9 [final concentration, 0.5 mg/mL], Tris-HCl buffer [pH 7.4; final concentration, 50 mmol/L], MgCl_2_ [final concentration, 0.5 mg/mL], alamethicin [final concentration, 0.25 µg/mL] and CIT [final concentration, 1.5 µg/mL]) was pre-incubated at 37 °C for 5 min. The total volume was then brought to 1 mL by adding UDPGA (final concentration, 3 mmol/L), and incubation was started at 37 °C. A 100-µL aliquot of the sample was collected from each mixture after 30, 60 and 240 min from the start of incubation. An equal amount of acetonitrile was then added to terminate the reaction. After centrifugation at 9000 *g* for 5 min at 4 °C, the supernatant was dried with nitrogen gas. Samples were stored at −30 °C until the analysis.

#### Quantification of CIT and detection of metabolites

Dried samples from the administration and permeability studies were re-dissolved in methanol for the analysis. The quantification of CIT in samples was conducted under the following analytical conditions: LC was performed using the Agilent 1290 Infinity LC System (Agilent Technology Ltd., Santa Clara, CA, USA), and separation was performed using a ZORBAX Eclipse plusC18 (100 mm, 2.1 mm, 1.8 µm; Agilent Technology Ltd.). The mobile phases used were 5 mmol/L acetic ammonium and methanol, and the solvent composition was increased in a linear gradient from 50% organic modifier to 85% at 7 min. The flow rate was 0.25 mL/min, the column oven temperature was 40 °C, and the injection volume was kept at 2 µL (administration study) or 0.1 µL (permeability study). MS was performed using the Agilent 6470 Triple Quadrupole LC/MS system (Agilent Technology, Ltd.). The ion source was the Agilent Jet Stream (AJS) (Positive/Negative mode), and the drying gas temperature and flow rate were 250 °C and 10 L/min, respectively, while the sheath gas temperature and flow rate were 400 °C and 12 L/min, respectively. The fragmentor voltage was 140 V, and the nozzle voltage was 1000 V. MRM transition was performed at m/z = 403 > 139 (30 eV), 297 (15 eV).

To detect metabolites by incubation using S9 without UDPGA, dried samples were re-dissolved in methanol. The LC system and analytical column were the same as described above. The mobile phases used were 0.1% formic acid and methanol, and the solvent composition was increased in a linear gradient from 10% organic modifier to 100% at 30 min. The flow rate was 0.2 mL/min. The quantification of CIT and search for metabolites of CIT were performed using an Agilent 6545 quadrupole time-of-flight mass spectrometer (Q-TOF) LC/MS system (Agilent Technologies, Ltd.). The drying gas temperature and fragmentor voltage were 350 °C and 120 V, respectively, and the other conditions were as described above.

To detect CIT glucuronide by incubation using S9 with UDPGA, dried samples were re-dissolved in acetonitrile. MS was performed using the Agilent 6530 Q-TOF LC/MS system (Agilent Technology, Ltd.). The mobile phases used were 5 mmol/L acetic ammonium and methanol, and the solvent composition was increased in a linear gradient from 10% organic modifier to 100% at 30 min. The flow rate was 0.2 mL/min, the column temperature was 40 °C, and the injection volume was 3 µL. the ion source was the AJS (Positive mode). Other conditions were as described above.

#### Statistical analyses

In incubation with S9, the CIT concentration and metabolite generation rate in humans and swine were analyzed using Student’s *t*-test or Welch’s test. p < 0.05 was accepted as indicating a significant difference. Statistical analyses were performed using the R version 3.5.0 (2018-04-23) (R Core Team (2018). R: A language and environment for statistical computing. R Foundation for Statistical Computing, Vienna, Austria. URL https://www.R-project.org/.).

## Results

### Toxicokinetics of CIT in swine

None of the swine administered CIT in IV or PO showed adverse clinical signs. Following IV dosing, the CIT concentration profile in plasma was fitted into a two-compartment model [16], as shown in Fig. 2A. The two-compartment model was described as follows:

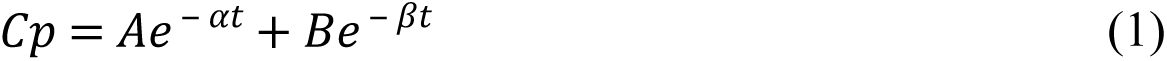

where Cp is the CIT concentration at time t (h) after IV. A and B are the initial concentrations at t=0 of the first and second phases of CIT concentration disposition, respectively, and α and β are the constants of the distribution and elimination phases, respectively. The toxicokinetic parameters were calculated using the equation above. The mean half-life of the distribution phase was 2.7 ± 3.6 h, and that of the terminal elimination phase was 17.7 ± 3.3 h (Table 1), showing that CIT seemed to be relative slow half-life in both phases. The volume of distribution (Vd) was greater than the total body water (1.005 ± 0.195 L/kg), and the total plasma clearance (Cltot) of CIT was estimated to 0.067 ± 0.019 L/h/kg. To calculate the bioavailability of CIT based on the different estimated parameters, the AUC was determined using the linear trapezoidal rule (1526 ± 336 ng·h/mL), with extrapolation to infinity.

**Fig 2.**
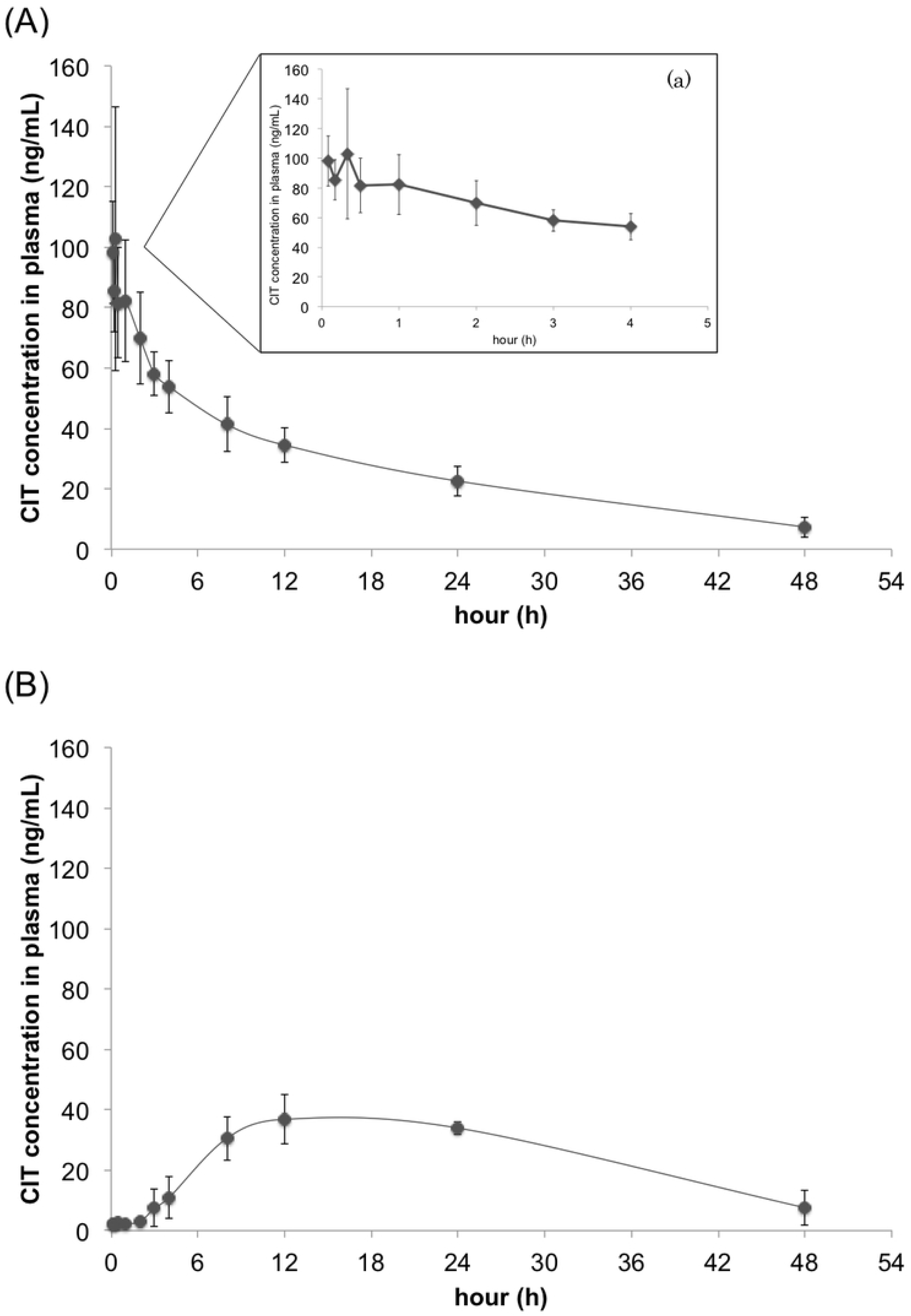
CIT concentration-time profiles in plasma after IV and PO administration to swine. IV (A) and PO administration (B) were performed with CIT (0.1 mg/kg·BW). The expanded time course (a) was showed in order to clarify to the profile from 5 min to 4 h. Control plasma was obtained on the day before administration. Values are presented as the mean ± standard deviation (SD). n=4 in both groups.

**Table 1.** Toxicokinetic parameters of CIT estimated from the two-compartment model in the plasma of swine following IV administration.

| Parameters | Route (IV) |
| --- | --- |
| BW (kg) | 9.35 ± 1.26 |
| A (ng/mL) | 45.6 ± 11.4 |
| $\alpha$ (h<sup>-1</sup>) | 0.718 ± 0.573 |
| B (ng/mL) | 56.4 ± 10.8 |
| $\beta$ (h<sup>-1</sup>) | 0.040 ± 0.008 |
| T1/2α (h) | 2.7 ± 3.6 |
| T1/2β (h) | 17.7 ± 3.3 |
| Vd (L/kg) | 1.005 ± 0.195 |
| Cl<sub>tot</sub> (L/h/kg) | 0.067 ± 0.019 |
| AUC (ng·h/mL) | 1526 ± 336 |

BW, body weight; C<sub>0</sub>, extrapolated plasma concentration at time zero; T<sub>1/2α</sub> and T<sub>1/2β</sub>, biological half-life of the distribution and the elimination; V<sub>d</sub>, apparent volume of distribution; Cl<sub>tot</sub>, plasma clearance; AUC, area under the curve, calculated using the linear trapezoidal rule from the curve extrapolated to infinity.
Values are presented as the mean ± SD.

The CIT concentration profile in plasma after PO administration is shown in Fig. 2B. The toxicokinetic parameters of PO administration were analyzed by a non-compartmental analysis [16]. The peak plasma concentration (Cmax, 38.2 ± 6.7 ng/mL) was observed between 15 ± 6 h (Tmax) after administration (Table 2). The mean residence time (MRT) was obtained from equation (2) as follows:

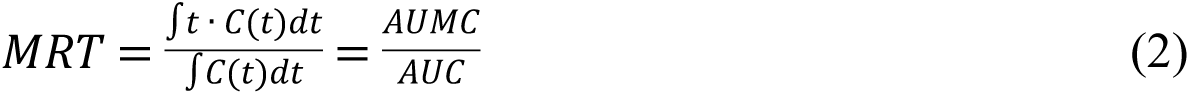

**Table 2.**
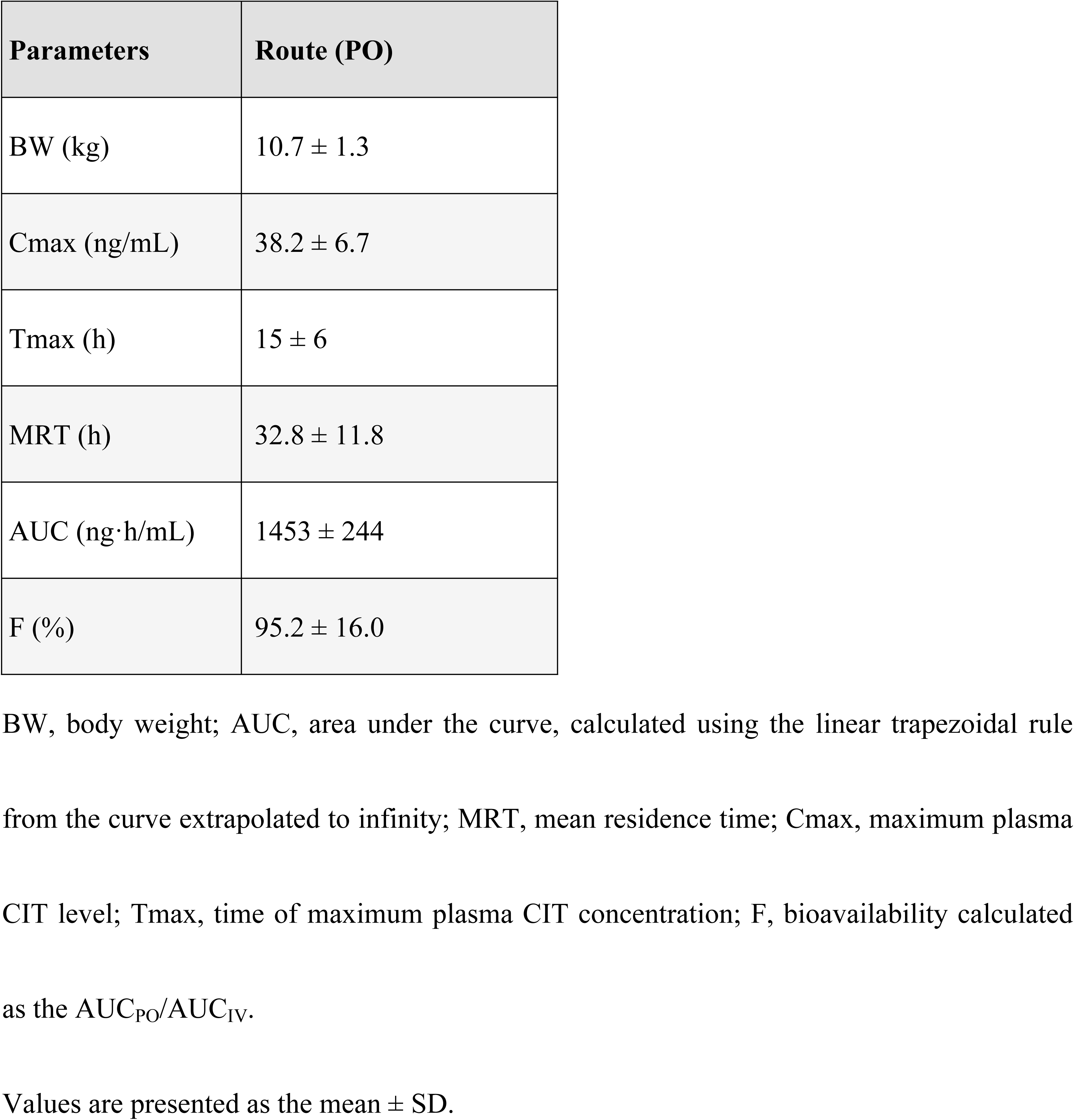
Toxicokinetic parameters of CIT estimated from the non-compartment model in the plasma of swine following PO administration.

| Parameters | Route (PO) |
| --- | --- |
| BW (kg) | $10.7 \pm 1.3$ |
| C<sub>max</sub> (ng/mL) | $38.2 \pm 6.7$ |
| T<sub>max</sub> (h) | $15 \pm 6$ |
| MRT (h) | $32.8 \pm 11.8$ |
| AUC (ng·h/mL) | $1453 \pm 244$ |
| F (%) | $95.2 \pm 16.0$ |

BW, body weight; AUC, area under the curve, calculated using the linear trapezoidal rule from the curve extrapolated to infinity; MRT, mean residence time; C<sub>max</sub>, maximum plasma CIT level; T<sub>max</sub>, time of maximum plasma CIT concentration; F, bioavailability calculated as the $AUC_{PO}/AUC_{IV}$ .
Values are presented as the mean ± SD.

The area under the first moment curve (AUMC) and AUC of PO and IV data were calculated using the trapezoidal rule. The MRT obtained from those values was relatively long (32.8 ± 11.8 h) (Table 2), demonstrating that CIT seemed to persist in the bodies of swine. For pigs, the bioavailability was estimated to be 95.2% ± 16.0%.

### Permeability study using Caco-2 cells

The results of the administration study showed that CIT had a high bioavailability in swine. In order to compare the intestinal permeability of CIT in humans with the bioavailability of CIT in swine, the permeability was investigated using Caco-2 cells. The Caco-2 cell model is an *in vitro* model used to evaluate the intestinal permeability and influence on the intestinal barrier function of chemical compounds in humans.

The apparent permeability coefficient (Papp) estimated from a Caco-2 permeability assay has been reported to correlate well with the human *in vivo* absorption data for many agents [19][20]. The TEER value is generally accepted to reflect the integrity of tight junction dynamics in Caco-2 cells [21], so the TEER was measured at 1 and 2 h after exposure to 3 and 10 µmol/L of CIT. Our results showed no marked change in the TEER over time at any concentration (data not shown). The transport rate of CIT from the AP to BL was calculated based on the concentration of CIT in the BL compartment. The Papp was calculated as described in a previous paper [17]. The Papp at 3 and 10 µmol/L CIT after incubation for 2 h was 52.2 × 10^−6^ and 42.6 × 10^−6^ (cm/s) in the AP-BL direction, respectively (Table 3). These findings indicated that human intestine cells were highly permeable to CIT *in vitro*.

**Table 3.** Papp at each CIT concentration.

| Parameter | 3 $\mu\text{mol/L}$ | 10 $\mu\text{mol/L}$ |
| --- | --- | --- |
| Papp ($\times 10^{-6}$ cm/s) | 52.2±28.3 | 42.6±17.7 |

Papp, apparent permeability coefficient of Caco-2 cells treated with 3 and 10 $\mu\text{mol/L}$ CIT in the AP chamber.
Values are presented as the mean ± SD.

### CIT elimination and the CIT metabolites profile following incubation with S9 *in vitro*

The fact that it took more than 40 h to eliminate CIT in plasma after IV and PO administration in swine *in vivo* experiments suggested that CIT was metabolized poorly in the liver and other organs. Therefore, to confirm the CIT metabolic activity and its metabolites produced in the liver and intestine, the residual CIT concentration after incubation *in vitro* with hepatic or intestinal S9 from swine was measured, and the profile of CIT metabolites produced was identified by using Q-TOF. These data obtained from the swine S9 fraction were subsequently compared with those obtained using human S9.

The elimination of CIT as well as the metabolites produced when CIT was incubated with S9 fractions supplemented with NADP was investigated. NADP is a coenzyme of dehydrogenase and used in metabolism assays with S9 popularly [22][23]. Incubation was conducted according to the method of Wu et al. [18]. As a result, hydroxylation, methylation (m/z = 432.2148), desaturation (m/z = 400.1886) and dihydroxylation (m/z = 434.1941) derivatives were detected as the main metabolites of CIT. The extracted ion chromatogram (EIC) and accurate mass spectrum of these metabolites are shown in Fig S1. These detected metabolites were confirmed from each product ion spectrum (data not shown). Although the concentration of CIT incubated with human hepatic S9 decreased with the duration of incubation, the concentration of CIT incubated with swine hepatic S9 was almost unchanged from 30 to 240 min of incubation (Fig. 3A). Furthermore, the CIT concentration incubated with hepatic S9 at 240 min was significantly lower with the human hepatic S9 than with the swine hepatic S9.

**Fig 3.**
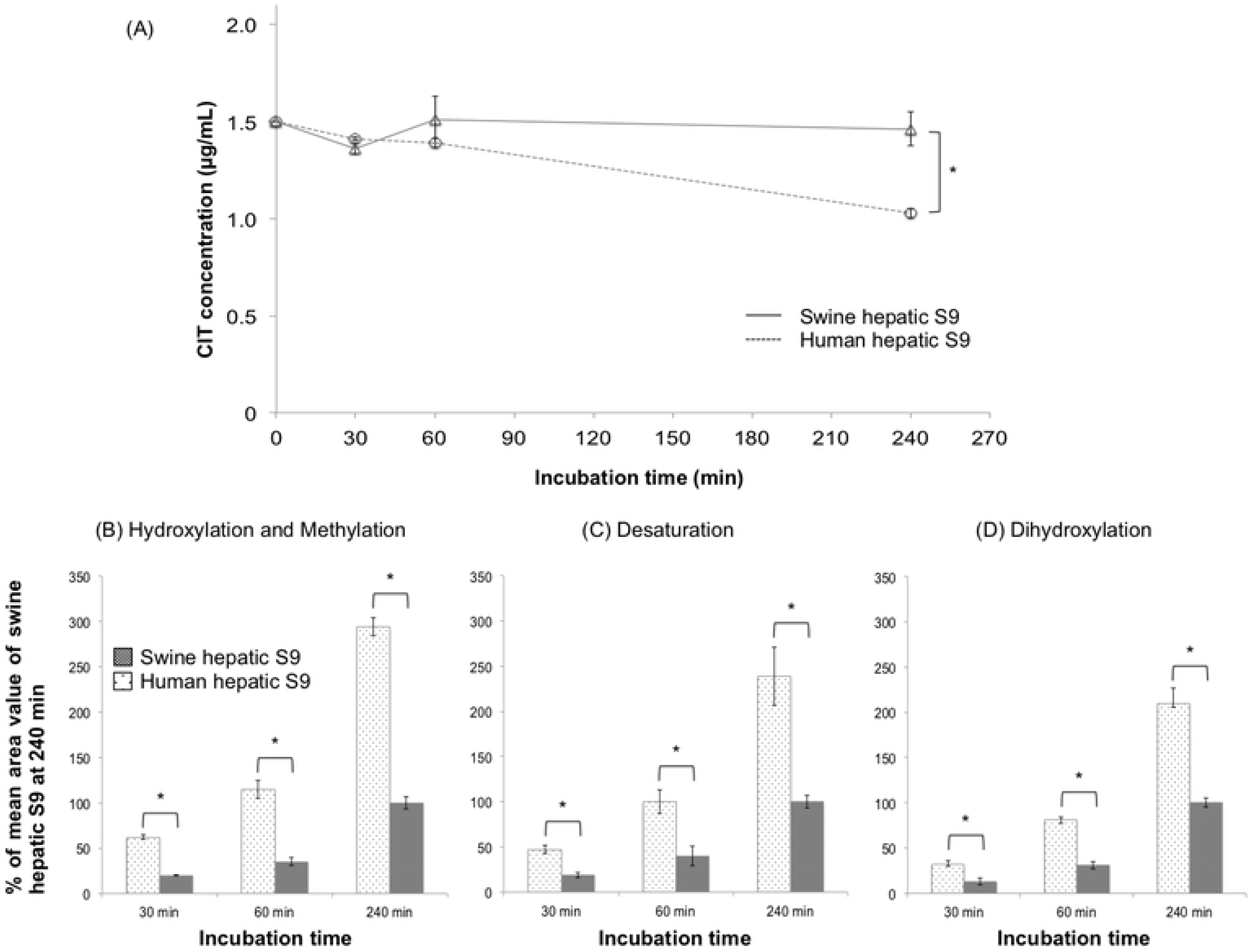
CIT concentration-time curves and the comparison of the main metabolites in humans and swine, obtained by incubating with the hepatic S9 fractions of humans and swine. CIT (1.5 µg/mL) was incubated with the hepatic S9 fractions of humans and swine supplemented with NADP as a coenzyme. The CIT concentrations up to 240 min after the start of incubation are described (A). The main metabolites produced in humans and swine are shown in (B), (C) and (D). Comparison of metabolites in humans and swine at each time was conducted with the mean area of each metabolite in swine as 100% at 240 minutes. (B), (C) and (D) show the percentage of hydroxylation and methylation (m/z=432.2148), desaturation (m/z=400.1886) and dihydroxylation (m/z=434.1941) derivatives by incubation with human and swine hepatic S9 respectively. Values are presented as the mean ± SD. Asterisks indicate a significant difference (p<0.05).

The main metabolites could not be quantified because standard substances of the metabolites detected were not commercially available. Therefore, comparison of metabolites in humans and swine at each time was conducted with the mean area of each metabolite in swine as 100% at 240 minutes. The rate of metabolite peak area by incubation with the human hepatic S9 fraction was two to three times higher than with the swine hepatic S9 fraction for all metabolites (Fig. 3B-D). These results indicate that human hepatic S9 has a greater ability to metabolize CIT than swine hepatic S9.

The concentration of CIT hardly decrease up to 240 min after incubation with the human intestinal S9 fraction, whereas it increased from 30 to 60 min after incubation with the swine intestinal S9 fraction, followed by almost no change from 60 to 240 min (Fig. 4A). Although the reason that the increase observed in incubation with swine intestinal S9 fraction was unclear, these indicated that CIT was hardly metabolized either intestinal S9 fractions. For the comparison, the same method as the comparison in S9 fractions supplemented with NADP was used. The rate of metabolite peak area by incubation with the human intestinal S9 fraction was roughly three times higher than with the swine intestinal S9 fraction for all metabolites (Fig. 4B-D). These were almost the same rates as in the above incubation using hepatic S9.

**Fig 4.**
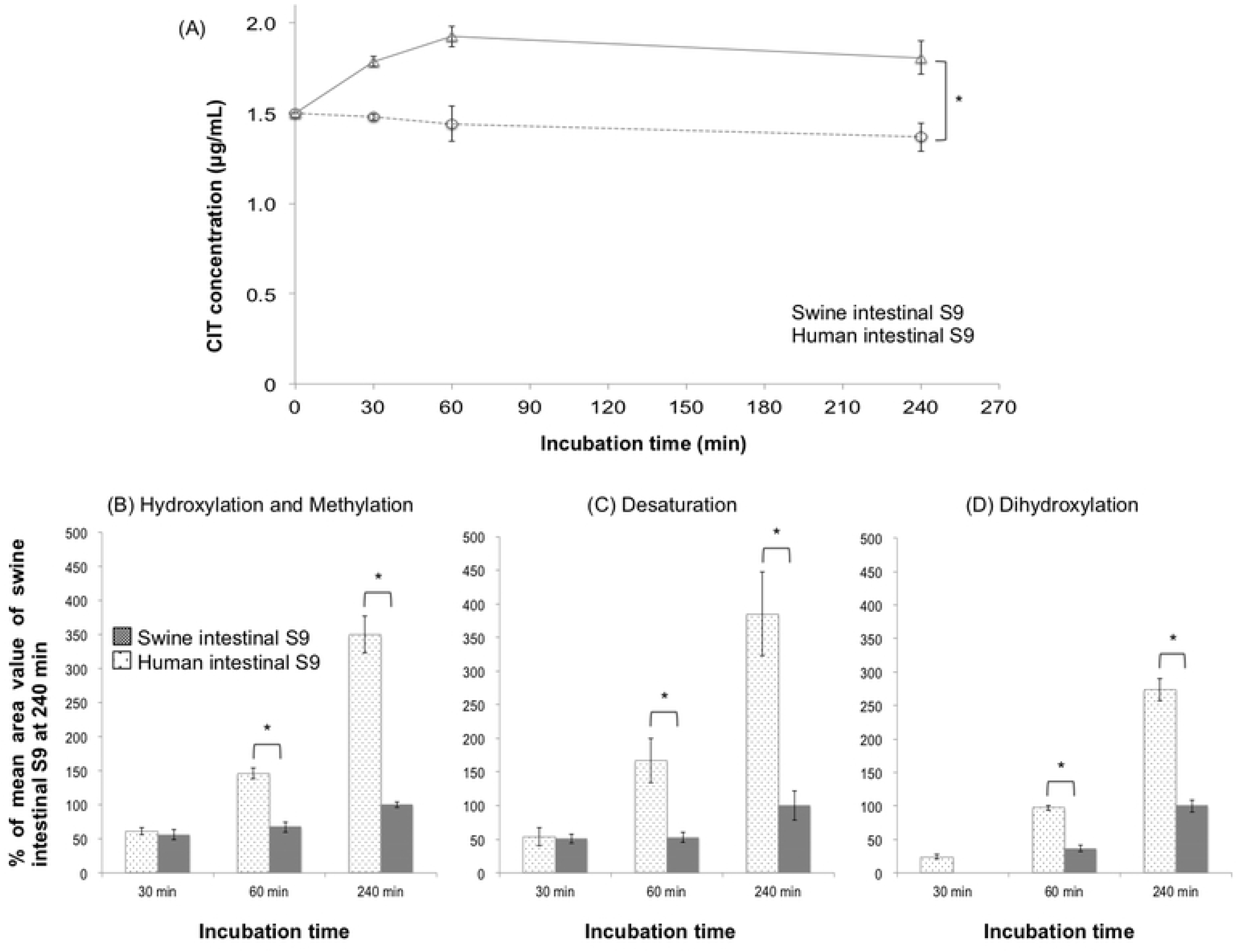
CIT concentration-time curves and the comparison of the main metabolites in humans and swine, obtained by incubating with the intestinal S9 fractions of humans and swine. CIT (1.5 µg/mL) was incubated with the intestinal S9 fractions of humans and swine supplemented with NADP. The CIT concentrations up to 240 min after incubation are described (A). The main metabolites produced in humans and swine are shown in (B), (C) and (D). Comparison of metabolites in humans and swine at each time was conducted with the mean area of each metabolite in swine as 100% at 240 minutes. (B), (C) and (D) show the percentage of hydroxylation and methylation (m/z=432.2148), desaturation (m/z=400.1886) and dihydroxylation (m/z=434.1941) derivatives by incubation with human and swine intestinal S9 respectively. Values are presented as the mean ± SD. Asterisks indicate a significant difference (p<0.05).

The peak areas of CIT metabolites produced by incubation with intestinal S9 were about one-third the size of those with hepatic S9 in both swine and humans (data not shown). However, these findings showed that CIT was hardly metabolized at all in the intestine of swine and humans.

Glucuronide is a well-known metabolite produced on detoxification of mycotoxins. However, following the above-mentioned incubation using S9 fractions with NADP as a coenzyme, no glucuronide was recognized in our swine or human metabolite models. Therefore, to compare the detoxication ability between swine and humans, the glucuronidation of CIT was examined under UDPGA conditions. UDPGA is a co-substrate used in the glucuronidation reaction [24]. Swine and human hepatic S9 were used for incubation, and CIT glucuronide was observed at 30, 60 and 240 min after incubation. The EIC and accurate mass spectrum of CIT glucuronide are shown in Fig. S2. Of note, with swine hepatic S9, CIT glucuronide was detected at 60 min and increased over time until 240 min, while CIT glucuronide was not detected through 240 min with human hepatic S9 (Fig. 5). CIT glucuronide in metabolite with intestinal S9 was not detected from either human or swine (data not shown).

**Fig 5.**
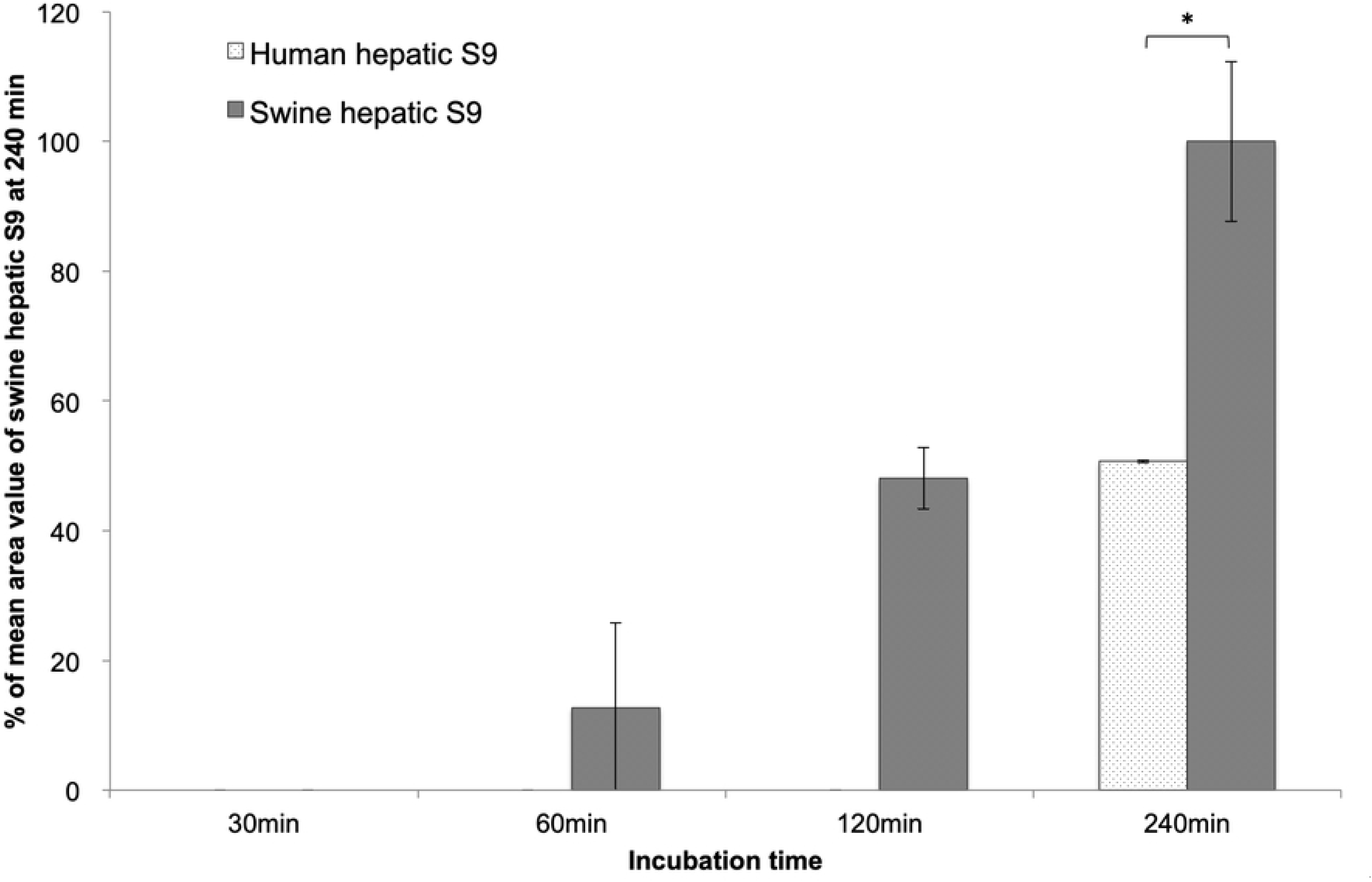
A comparison of CIT glucuronidation during incubation with human and swine S9 fractions supplemented with UDPGA. CIT (1.5 µg/mL) was incubated with human and swine hepatic S9 fractions supplemented with UDPGA as a coenzyme. CIT glucuronide at 30, 60 and 240 min was measured by Q-TOF. Comparison of metabolites in humans and swine at each time was conducted with the mean area of each metabolite in swine as 100% at 240 minutes. Asterisks indicate a significant difference.

Because the standard substance of CIT glucuronide was not commercially available, for the comparison, the same method as the comparison in S9 fractions supplemented with NADP was used. The rate of the mean area of CIT glucuronide at 240 min of incubation with swine hepatic S9 was about twice that with human hepatic S9 (Fig. 5). The CIT concentration decreased when using both human and swine hepatic S9 with UDPGA as coenzyme (Fig. 6). Furthermore, at 240 min of incubation, the CIT concentration with human hepatic S9 with UDPGA was significantly lower than that with swine S9 (Fig. 6). This showed that the CIT metabolism was greater in humans than in swine, although CIT glucuronidation hardly occurred at all in humans.

**Fig 6.**
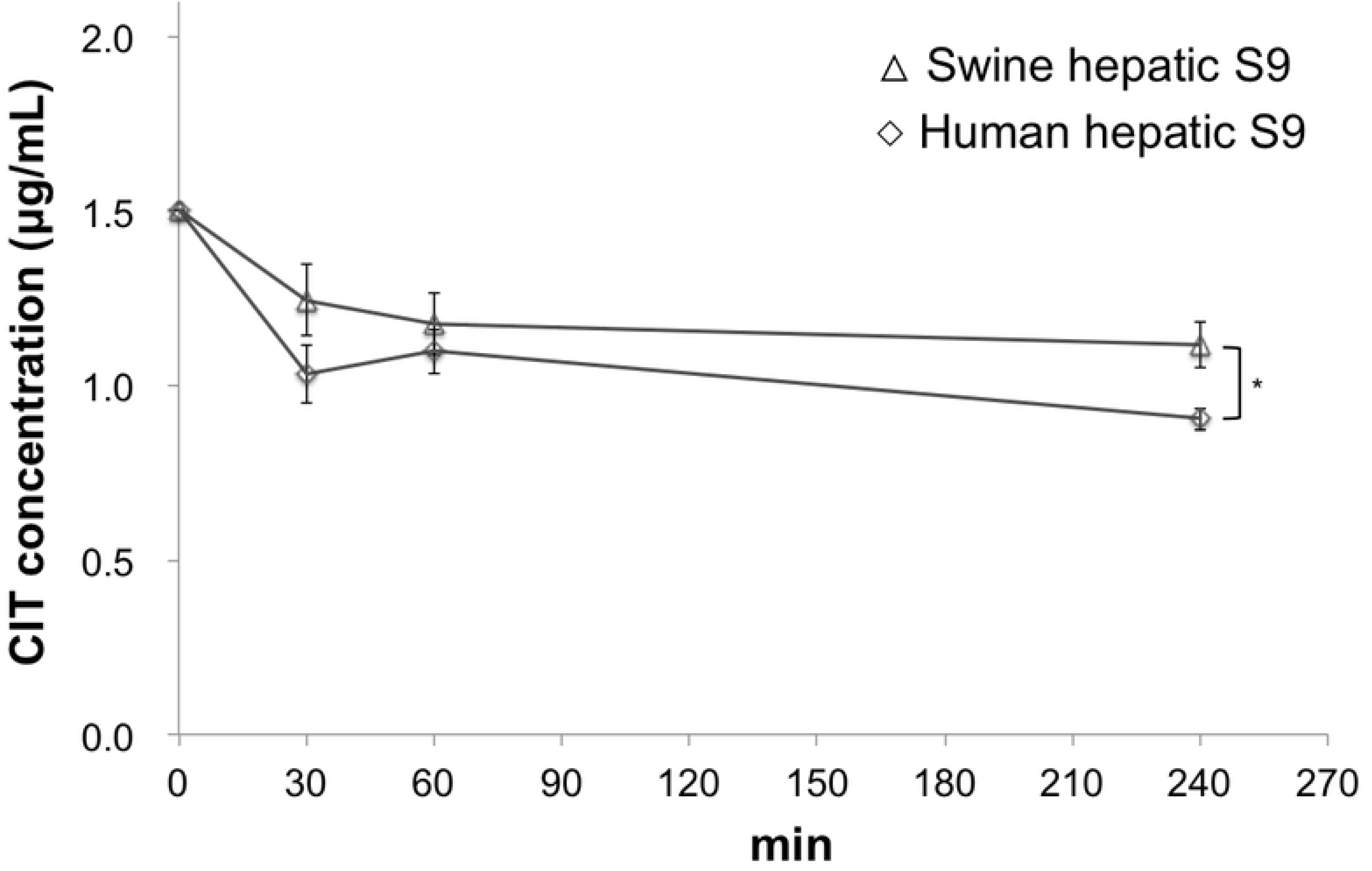
CIT concentrations–time curves incubated with S9 fractions supplemented with UDPGA. CIT (1.5 µg/mL) was incubated with human and swine hepatic S9 containing UDPGA. The CIT concentrations up to 240 min after incubation are described. Values are presented as the mean ± SD. Asterisks indicate a significant difference (p<0.05).

## Discussion

Swine are known to share some similarities with humans physiologically and anatomically, including food habits. The toxicokinetics of CIT were therefore investigated in swine for extrapolation to humans. The bioavailability of mycotoxins has been reported for several major compounds. Deoxynivalenol (48%-65% in swine) [25] [26] and ochratoxin A (65.7% in swine) [27] have a relatively high bioavailability, and zearalenone was shown to be absorbed at 80%–85% in the bodies of swine [28]. The results of the present study suggested that CIT was also a mycotoxin with a high bioavailability similar to these compounds (Table 2). Generally, fat-soluble substances appear to be easily absorbed by the intestine through passive transport, which may be one reason CIT showed a high bioavailability.

The CIT concentration in plasma showed almost no increase until three hours after its oral administration to swine (Fig. 2B). The dwell time of digesta in the stomach of swine has been reported to range from one to three hours [29]. As CIT was administered with feed in this study, CIT may have been retained with digesta in the stomach. A marked increase in the plasma CIT concentration in the present study was noted from three hours after administration, suggesting that it was poorly absorbed from the stomach.

The CIT concentration-time curve derived from the IV administration showed that a second peak appeared at two hours after administration (Fig. 2A), possibly due to enterohepatic circulation. Zearalenone is a mycotoxin that undergoes enterohepatic circulation, and a second peak has been observed in the plasma following IV administration in swine [30]. Biehl et al. further showed that the second peak of zearalenone disappeared when the bile was removed, prompting them to conclude that the peak indicated enterohepatic circulation [28]. Similarly, the second peak of the curve observed in this study may indicate enterohepatic circulation. Glucuronic acid conjugate is generally excreted into the bile and is considered relatively susceptible to enterohepatic circulation. However, CIT glucuronide was produced slowly in our *in vitro* study using S9 and was not detected at all in the plasma following administration *in vivo*. As other conjugates were not investigated in this study, future studies should explore the enterohepatic circulation of CIT.

In the present study, out of concern for the animals’ welfare, frequent blood drawing within the first hour after administration was performed under anesthesia. While we cannot discount the effects of anesthesia, the elimination of CIT from the body of swine appeared to be quite slow. Ueno et al. reported the distribution and elimination of CIT in their *in vivo* study in 1972 [11]. After administering extracted CIT subcutaneously to rats, they noted that CIT was rapidly distributed to the main organs, such as the liver, kidney and heart, from the administration site. They also found that the highest concentration of CIT was in the liver after 8 h, and <1% of the total dose administered could still be detected in the liver even after 52 h. In the present study, the Vd of CIT in swine was greater than 1 L/kg (Table 1). Generally, drugs with a Vd exceeding 1 L/kg are considered to be widely distributed to the body tissue [31]. Therefore, CIT is suspected to be distributed widely to the body tissue. Regarding other mycotoxins, ochratoxin A (84.5 h in swine, 840 h in monkey) [27][32] and aflatoxin B1 (91.8 h in rat) [33] reportedly have long elimination half-lives. In contrast, the elimination half-lives of fumonisin B1 (182 min in swine) and deoxynivalenol (7.2-15.2 h in swine) are reported to be short [34]. In this respect, CIT is a mycotoxin with a relatively long half-life (Table 1). Hou et al. proposed that CIT bound plasma albumin [35]. Ochratoxin A is generally accepted to bind plasma albumin [36][37], which may explain why CIT showed a long elimination half-life. In addition, Uraguchi reported that extract from yellow rice had a lethal effect, even when exposed rats were given daily doses of 1/300 of LD_50_ PO for 16 months. In the present study, the toxicokinetics of CIT in swine showed that CIT had a high bioavailability and persisted in the body for a relatively long period of time. This suggested that CIT might accumulate in the body with chronic exposure, and this result was considered to reflect the above report describing the adverse effects of chronic exposure.

To estimate the bioavailability in humans, the metabolites in the S9 fraction of the liver and intestine of humans were examined and compared with those in swine in an *in vitro* experiment. Regarding the metabolites present in the S9 fraction of humans, the main metabolites, such as the hydroxylation and methylation, desaturation and dihydroxylation derivatives, were the same as those in the S9 fraction in swine (Figs. 3 and 4). The metabolite-producing ability of the intestinal S9 fraction was about one-third of that in the hepatic S9 fraction in both species (data not shown), and the CIT concentration had hardly been reduced at all, even after 240 min. This suggested that CIT would be hardly metabolized in the intestine of swine and humans. In contrast, marked differences were noted in the metabolite-producing ability of the hepatic S9 fraction, with the metabolization being higher when using human S9 than when using swine S9. Although glucuronide was detected among the metabolites produced by the hepatic S9 fractions of both species supplemented with UDPGA, the glucuronidation of CIT was shown to be slower in humans than in swine (Fig. 5). However, the CIT concentration was significantly lower when using human hepatic S9 than when using swine hepatic S9 (Fig. 6). Interspecies differences in the hepatic glucuronidation of deoxynivalenol have been reported [38]. Similarly, there may have also been interspecies differences in the glucuronidation of CIT, which may be one reason that the glucuronidation rate of CIT was slower in the human hepatic S9 fraction than in the swine hepatic S9 fraction (Fig 5). Overall, these results of our *in vitro* study using S9 suggested that the CIT metabolism in the liver would be faster in humans than in swine (Fig. 3 and 6).

The permeability was also examined using Caco-2 cells. In this study, CIT showed a high Papp (Table 3) despite no marked effect on the TEER. Furthermore, this value was higher than the Papp of deoxynivalenol (0.19 ± 0.02 [×10^−6^ cm/min]) [39] and zearalenone (10.4 ± 4.7 [×10^−6^ cm/s]) [40], although it was lower than that of aflatoxin M1 (105.10 ± 7.98 [×10^−6^ cm/s]) [41]. The permeability coefficient from the Caco-2 cell assay was shown to correlate with the bioavailability and intestinal absorbency following a sigmoidal curve, wherein a substance with a high permeability coefficient had a high absorbency [19] [20]. Considering that the results of our *in vivo* study using swine (Table 2), indicated relatively higher bioavailability of CIT, the result of Caco-2 study suggested that CIT would have a similarly high bioavailability in humans as in swine.

In conclusion, the present study explored the toxicokinetics of CIT through an *in vivo* study using swine and predicted the characteristics of CIT in humans through a comparison *in vitro* study using Caco-2 cells and S9 fractions. The results indicated that CIT had a high bioavailability in swine and persisted in the body for a relatively long time. Therefore, CIT may be brought adverse effects in the context of cumulative by chronic exposure. In addition, a comparison of findings in humans and swine *in vitro* suggested that CIT had a similarly high bioavailability in humans as in swine, although CIT fortunately appears to be metabolized more quickly in humans than in swine.

## Acknowledgments

This work was supported by the Health and Labour Sciences Research Grants (Research on Food Safety, H28-shokuhin-ippan-004) from the Ministry of Health, Labour and Welfare of Japan. We thank Yoshihito Shimazu, Laboratory of Food and Physiological Sciences, Department of Life and Food Sciences, School of Life and Environmental Sciences, Azabu University and Masashi Sekimoto, Laboratory of Environmental Hygiene, Department of Environmental Science, School of Life and Environmental Science, Azabu University, for cooperating with us to prepare swine intestinal S9.

## Supporting information

**S1 Fig. The extracted ion chromatogram (EIC) and mass spectrum of the main metabolites of CIT incubated with S9.** The EIC and mass spectrum were analyzed by Q-TOF. The compounds presumably produced by incubation with S9 fractions supplemented with NADP were as follows: A, B and C represent the EIC and mass spectra of hydroxylation and methylation (EIC 433.22000), desaturation (EIC 401.19000) and dihydroxylation (EIC 435.20000), respectively, at 240 min after incubation with swine hepatic S9.

**S2 Fig. The EIC and mass spectrum of CIT glucuronide generated by incubation with S9 including UDPGA.** The EIC and mass spectrum of CIT glucuronide produced by incubation with swine hepatic S9 supplemented with UDPGA were analyzed by Q-TOF. The EIC and mass spectrum of CIT glucuronide as typical are showed those detected at 240 min after incubation with swine hepatic S9.

